# Modular preprocessing pipelines can reintroduce artifacts into fMRI data

**DOI:** 10.1101/407676

**Authors:** Martin A. Lindquist, Stephan Geuter, Tor D. Wager, Brian S. Caffo

## Abstract

The preprocessing pipelines typically used in both task and restingstate fMRI (rs-fMRI) analysis are modular in nature: They are composed of a number of separate filtering/regression steps, including removal of head motion covariates and band-pass filtering, performed sequentially and in a flexible order. In this paper we illustrate the shortcomings of this approach, as we show how later preprocessing steps can reintroduce artifacts previously removed from the data in prior preprocessing steps. We show that each regression step is a geometric projection of data onto a subspace, and that performing a sequence of projections can move the data into subspaces no longer orthogonal to those previously removed, reintroducing signal related to nuisance covariates. Thus, linear filtering operations are not commutative, and the order in which the preprocessing steps are performed is critical. These issues can arise in practice when any combination of standard preprocessing steps—including motion regression, scrubbing, component-based correction, global signal regression, and temporal filtering—are performed sequentially. In this work we focus primarily on rs-fMRI. We illustrate the problem both theoretically and empirically through application to a test-retest rs-fMRI data set, and suggest remedies. These include (a) combining all steps into a single linear filter, or (b) sequential orthogonalization of covariates/linear filters performed in series.

## 1 Introduction

In the past decade, resting-state functional magnetic resonance imaging (rs-fMRI) data has been increasingly used to study intrinsic functional connectivity in the human brain (Biswal et al., 1995). Using rs-fMRI it has been shown that fluctuations in the blood oxygen level dependent (BOLD) signal in spatially distant regions of the brain are strongly correlated (Beckmann et al., 2005; De Luca et al., 2006; Yeo et al., 2011). While the exact mechanisms driving these correlations remain unclear, it has been hypothesized that it may be due to fluctuations in spontaneous neural activity. Neuroscientists have become increasingly interested in studying the correlation between spontaneous BOLD signals from different brain regions in order to learn more about the inner workings of the brain (Van Den Heuvel and Pol, 2010).

The analysis is complicated by the fact that the measured BOLD signal consists of both changes induced by neuronal activation, as well as non-neuronal fluctuations. Here the former is the signal of interest, while the latter is considered nuisance signal. Examples of such non-neuronal fluctuations include drift, spiking artifacts, motion-related artifacts, and fluctuations due to physiological sources such as heart rate and respiration. Failure to properly control for these types of noise can have significant impact on subsequent analysis. For example, head motion has been shown to have systematic effects on resting state functional connectivity measures (Van Dijk et al., 2012; Power et al., 2014), and comparisons between groups of subjects with different levels of head motion have yielded difference maps that could be mistaken for interesting neuronal effects (Van Dijk et al., 2012). Furthermore, it has been shown there are significant correlation between changes in cardiac and respiratory rates and the BOLD signal (Birn et al., 2006; Shmueli et al., 2007; Wise et al., 2004). These nuisance fluctuations risk artificially inflating functional connectivity measures, or even creating spurious findings (Murphy et al., 2013).

In general, the relative influence of these different non-neuronal fluctuations depend on a number of factors (Caballero-Gaudes and Reynolds, 2017). However, it is clear that if not treated properly these sources of variation can induce spurious functional connectivity between different brain regions. Thus, it is of great interest to reduce their effects on the analysis of rs-fMRI data.

For these reasons, rs-fMRI data are subjected to a series of preprocessing steps (Weissenbacher et al., 2009; Yan and Zang, 2010; Caballero-Gaudes and Reynolds, 2017) prior to analysis. Typically, each step consists of a separate algorithm—often a linear regression of data on nuisance covariates, with residuals used in subsequent analysis, or a linear filtering operation—designed to remove a specific type or class of artifacts. These steps can include motion regression (Fox et al., 2009; Weissenbacher et al., 2009), scrubbing (Power et al., 2012, 2014) or spike regression (Lemieux et al., 2007; Satterthwaite et al., 2013), nuisance regression (e.g., the removal of signal from white matter (WM) and ventricular cerebrospinal fluid (CSF) tissues; Behzadi et al. (2007); Muschelli et al. (2014)), the removal of global signal (Fox et al., 2005), and temporal filtering including low-and high-pass filters (Biswal et al., 1995; Cordes et al., 2001). The most popular versions of all of these can be expressed as linear filtering operations.

There are a large variety of rs-fMRI pre-processing pipelines (i.e., combinations of such modular preprocessing steps) that vary in the specific operations they perform on the data, as well as the order in which they are performed. In fact, a paper by Carp (2012) showed that there are nearly as many unique analysis pipelines in the literature as there were studies^1^. While most of the preprocessing steps that are performed are essential, there is relatively little understanding of the effects they have on both the spatial and temporal correlation structure of the resulting data. Importantly, there is a general lack of knowledge regarding potential interactions among the individual preprocessing steps, and how the order in which they are performed impacts the resulting analysis. To date, there is no consensus standard of what steps should be included in a pipeline, or in which order they should be performed.

Whatever the procedures performed, most popular pipelines have gravitated towards removing nuisance covariates in a series of sequential steps, which simplifies the analysis, without considering the order in which they are performed. However, we show here that these linear filtering operations are not commutative, and furthermore, that performing them in series can re-introduce nuisance signal removed in previous steps.

### A geometric approach

When working with linear models it is often fruitful to take a geometric approach towards understanding their behavior by viewing them as linear projections. Here the fitted value in the regression is seen as the orthogonal projection of the data onto the subspace spanned by the columns of the design matrix. Similarly, the residuals are the projection onto the subspace that is orthogonal to the columns of the design matrix, and are thus uncorrelated with the nuisance components that make up these columns. This is illustrated graphically in Fig. 1, where the data (represented as an *n*-dimensional point) is projected onto the *p*-dimensional subspace spanned by the columns of the design matrix (shown in light blue).

**Figure 1:**
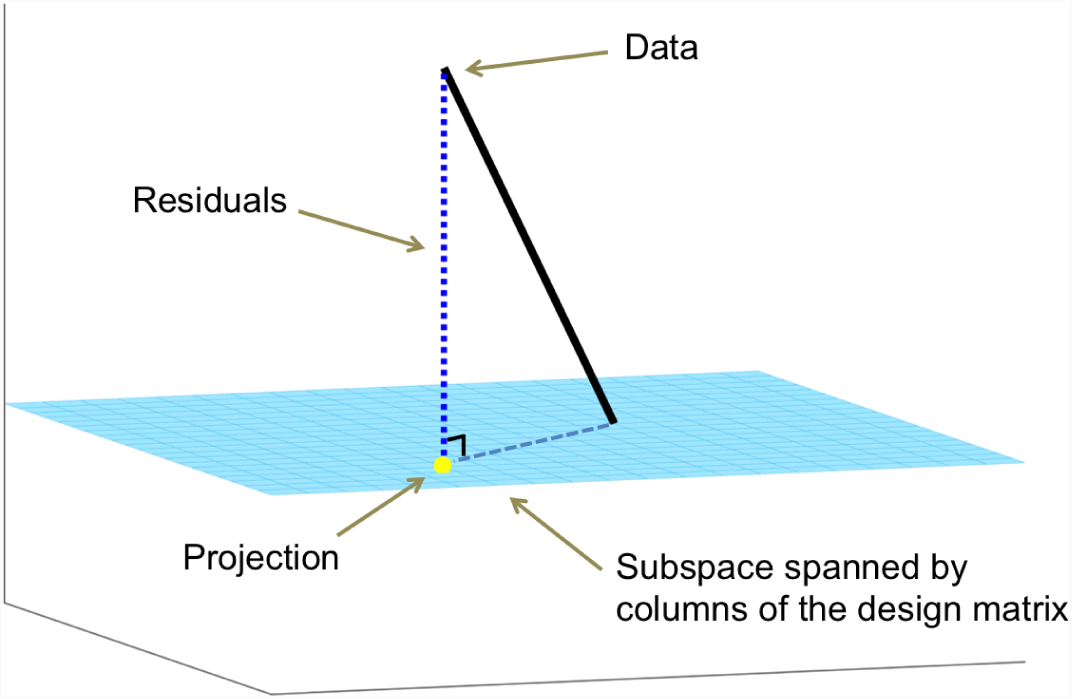
An illustration of the geometry of linear projections. The *n*-dimensional data is projected onto the *p*-dimensional subspace spanned by the columns of the design matrix (shown in light blue). The projection (shown in yellow) corresponds to the fitted value. The residuals (i.e., the data minus the fitted value) lie in a subspace that is orthogonal to the columns of the design matrix.

In this paper we take a geometric approach towards analyzing modular rs-fMRI preprocessing pipelines. We express commonly used processing steps, such as motion regression, spike regression, nuisance regression, and temporal filtering, as projections onto a subspace of the full *n*-dimensional space (where *n* represents the number of time points) in which the data resides. This allows us to evaluate how different processing steps interact with one another, and how they in many settings actually counteract one another. In particular, it allows us to illustrate that both the order and the manner in which preprocessing steps are performed is critically important for being able to properly interpret subsequent analysis. Using our geometrical approach we illustrate how, if not performed carefully, certain preprocessing techniques have the effect of re-introducing previously removed artifacts back into the signal.

We illustrate this issue primarily in the context of motion regression and temporal filtering, both theoretically and using test re-test rs-fMRI data. We show how performing high-pass filtering after motion regression reintroduces motion artifacts into the data. Similarly, performing motion regression after high-pass filtering reintroduces unwanted frequency components into the signal. The latter issue has been discussed in a number of papers, including work by Hallquist et al. (2013). Here we take their results and place it into a broader geometrical framework.

Although we focus on a few specific cases, these issues arise anytime multiple preprocessing steps are used that act as projections onto a subspace of the original data-space. In addition, though we focus on rs-fMRI data in this paper, we stress that these issues can potentially arise in task fMRI as well if modular preprocessing is performed. To avoid these issues we recommend that researchers either perform a simultaneous regression on all nuisance covariates, or alternatively orthogonalize later covariates with respect to the ones removed in earlier stages of the pipeline. This latter point implies one must orthogonalize both the data, and all subsequent projections, in order to maintain data orthogonality with the current projection. Interestingly, these approaches have become standard practice in the preprocessing of task-fMRI data.

We believe our framework has the potential to simplify the critical evaluation of preprocessing pipelines, and identify areas where problems can occur. In this paper, we illustrate that the issues discussed in this work can have significant effect on subsequent analysis, and we therefore urge that special care be taken when performing preprocessing on rs-fMRI data. We further highlight the need to critically revisit previous work on rs-fMRI data work that may not have adequately controlled for these types of effects.

## 2 Theoretical Background

In this section we show theoretically how the use of modular preprocessing steps can reintroduce artifacts that were removed in a previous step. We also provide recommendations for circumventing these issues.

### 2.1 A geometric approach for evaluating pipelines

The issue we discuss in this paper can potentially arise anytime one uses a preprocessing step that projects the data onto a subspace of the *n*-dimensional space in which it resides. This includes any technique that utilizes a linear model framework to remove artifacts from the signal. To illustrate the problem, let us consider the case of performing motion regression and temporal filtering sequentially. Hallquist et al. (2013) previously showed that in this setting nuisance-related variation can be reintroduced into frequencies that were previously suppressed by the filter. Here we generalize their findings to incorporate a wider array of preprocessing steps, and place it into a general mathematical framework.

Let **y** be an *n*-dimensional vector containing the rs-fMRI signal from a specific voxel in the brain. Further, let **X** be an *n* × *p* design matrix containing the nuisance regressors we seek to remove from **y**. In our illustration let us assume that *p* = 24, corresponding to (i) the six motion regressors obtained after rigid body transformation; (ii) the regressors squared; (iii) their first order difference; and (iv) the first order difference squared.

To remove the effects of these nuisance components, the next step is to regress them out of **y**. To do so, we fit a linear model on the form:

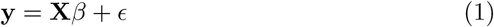

It is well known that the least-squares estimate of *β* is given by

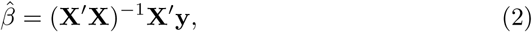

and the fitted value can be expressed as:

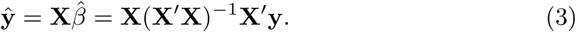

Here the term 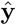 represents the estimated motion in the voxel and corresponds to the nuisance signal that we seek to remove from **y**.

Here it is useful to take a projection approach towards performing linear regression. To do so we define

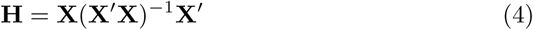

and thus we can write 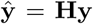 In this setting, **H** is referred to as a projection matrix and its application has the effect of projecting the data onto the space spanned by the columns of the design matrix **X**. Importantly, projection matrices are both idempotent (**H** = **H**^2^) and symmetric (**H** =**H**′).

To remove the effects of motion we subtract 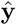 from the data and obtain the residual 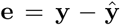 In terms of the projection matrix, we can write this as **e** = (**I** − **H**)**y**. This term is now our signal of interest. It is important to note that **P**_1_ = **I H** is also a projection matrix, and it projects the data onto a subspace that is orthogonal to the columns of the design matrix **X**. Thus, **e** resides in a subspace orthogonal to **X**. Hence,

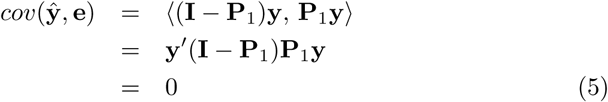

and the data is uncorrelated with the motion, as required. The last equality holds as 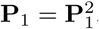 (Here ⟨.,.⟩ represents the Euclidean inner product.)

At this point we have, to the extent possible by linear projections, successfully removed the effects of the nuisance components that make up the columns of **X** from the data. However, it is important to realize that researchers at this stage typically perform additional modular preprocessing steps that act as projections of the data into new subspaces. For example, this would occur if we were to continue by next performing a preprocesing step such as spike regression, component-based correction (CompCor), global signal regression, or temporal filtering. This can have the unfortunate side-effect of projecting the data back into the space spanned by the motion regressors in **X**, and thus reintroducing the effects of components that had previously been removed. This will happen if the two sub-spaces are not orthogonal to each other.

To illustrate, suppose we perform high-pass filtering on the motion-regressed data. Filtering can be expressed in a linear model using a series of sine and cosine functions as regressors. Fitting this model projects the data onto a space orthogonal to the frequency components one seeks to remove. We can express this operation as **e**_*f*_ = **P**_2_**e**. Unfortunately, this operation projects the data into a subspace that is no longer constrained to be orthogonal to the space spanned by **X**. This has the effect of partially reintroducing a correlation between the motion and the signal being studied. This can be noted by observing that

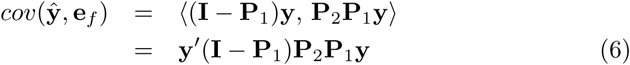

is no longer guaranteed to be zero, and indeed in most situations will in fact be non-zero. This illustrates the problem from a theoretical point of view.

Empirically, we illustrate the problem on real resting-state data in Section 3. Fig. 2D-F summarizes some of the results. Panel (D) shows the rs-fMRI time course and the estimated motion from a specific voxel of interest. The two time courses show a large positive correlation (*r* = 0.7622). After motion correction, the processed data and the estimated motion are now uncorrelated (*r* = 0), as would be expected from Eq. 5. Finally, after high-pass filtering the data and the estimated motion are now negatively correlated (*r* = −0.3458). Clearly, high-pass filtering has partially reintroduced a correlation between the data and motion.

**Figure 2:**
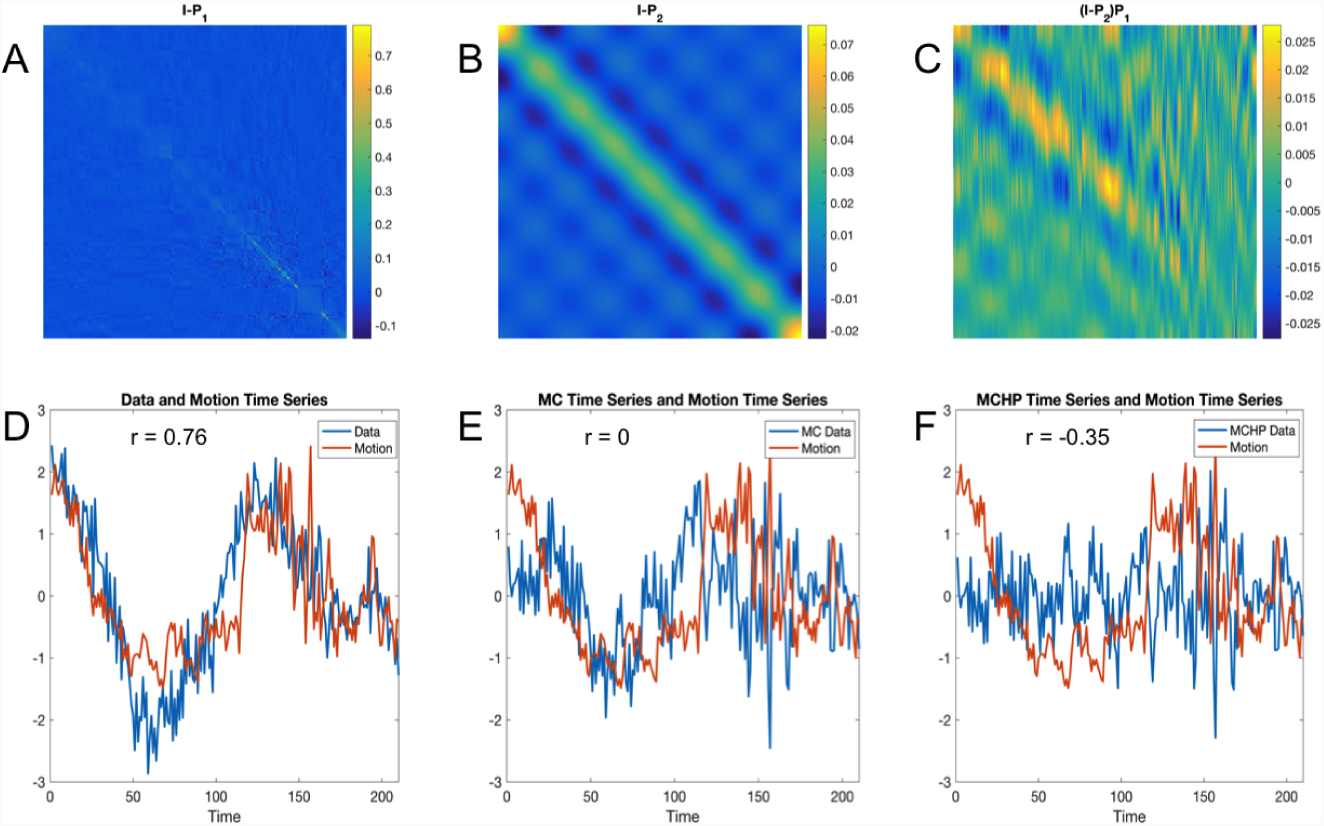
Examples of (A) a projection matrix **I** − **P**1, removing empirically computed motion artifacts; (B) a temporal filtering projection matrix **I** − **P**_2_, and (C) the product of both projection matrices (**I***-* **P**_2_)**P**_1_, which is clearly non-zero. (D) The rs-fMRI time course (blue) and the estimated motion (red) from a voxel of interest. The two time courses are positively correlated (*r* = 0.7622).(E) The same data after motion correction (blue) together with the estimated motion (red). The two time courses are uncorrelated (*r* = 0). (F) The same data in (E) after high-pass filtering (blue) together with the estimated motion (red). The two time courses are negatively correlated (*r* = 0.3458). Hence, high-pass filtering has reintroduced a correlation between the data and motion.

Eq. 6 highlights that a requirement for the data and motion to be uncorrelated is that (**I** − **P**_1_)**P**_2_**P**_1_ = **0**. Apart from trivial cases (e.g., **P**_1_ = **0**, **P**_2_ = **0**, **P**_1_**P**_2_ = **0**, or **P**_2_ = **AP**_1_ for some **A**), this holds if **P**_1_ = **P**_2_**P**_1_, or equivalently that **P**_1_(**I** − **P**_2_) = **0**. This implies that the filter projects the data onto the space orthogonal to that spanned by **X**. This will almost certainly not hold unless the filter is explicitly designed for this purpose, for example if the regressors forming **P**_2_ are orthogonalized with respect to those forming **P**_1_.

Empirically, this issue is illustrated in Fig. 2. Panels A-C show examples of the matrices **I** − **P**_1_, **I P**_2_, and (**I** − **P**_2_)**P**_1_, respectively, corresponding to motion regression, high-pass filtering, and their product. Clearly the last term is not 0, and hence the covariance between **e**_*f*_ and 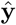 will not be equal to 0, as shown empirically in Panel F.

Note a related problem arises if temporal filtering is first applied, followed by motion regression. Here the later step reintroduces signal into frequency bands which had previously been removed. The reason is similar to that described above as the space orthogonal to **X** need not be orthogonal to the space spanned by terms corresponding to the retained frequency components. This issue was discussed in detail in Hallquist et al.(2013).

The issue is further illustrated in Fig. 3. Here we show graphically what happens when a point in *n*-dimensional space, represented by the black vector, is repeatedly projected onto different subspaces. Suppose the point is first projected onto the light-blue subspace. Note that though shown in 2-dimensions, this subspace is actually (*n – p*_1_)-dimensional, where *p*_1_ are the number of columns in the design matrix used to create the first projection matrix. The projection is shown in yellow. Further suppose, this point is subsequently projected onto the gray subspace. This subspace is (*n – p*_2_)-dimensional, where *p*_2_ are the number of columns in the design matrix used to create the second projection matrix. The new location is now shown in green. Here we note that in the plot to the left, where the two subspaces are orthogonal to one another, the point simultaneously lies in both subspaces. In the plot to the right, where the subspaces are no longer orthogonal, the point is not located in the light-blue subspace.

**Figure 3:**
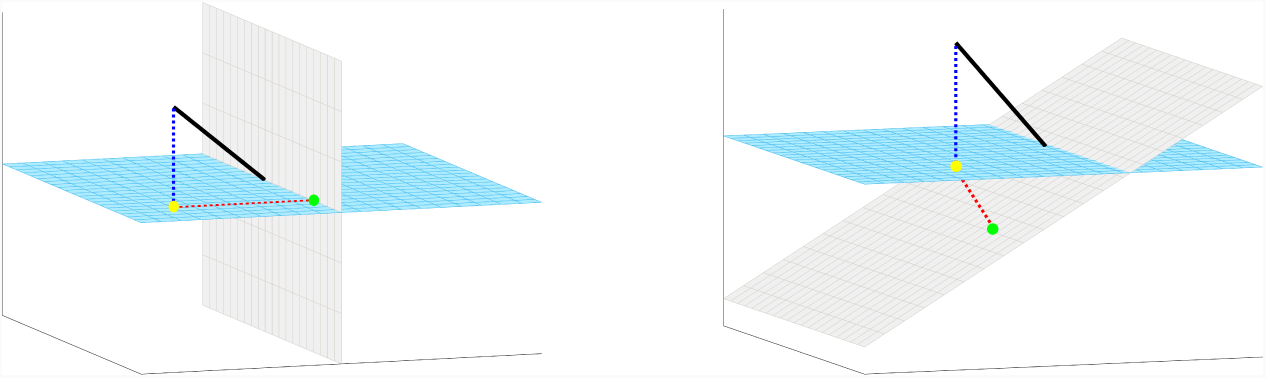
Illustration of the effect of multiple projections. Consider a point in *n*-dimensional space represented by the black vector. The point is first projected onto the light-blue subspace; see yellow point. The first projection is subsequently projected onto the gray subspace; see green point. In the plot to the left the two subspaces are orthogonal to one another and thus the second projection simultaneously lies in both subspaces. In the plot to the right the green point is no longer located in the light blue subspace and thus not orthogonal to the confounds defining that subspace.

To tie this example to the preprocessing setting, let us consider the light-blue subspace to be the space orthogonal to the motion regressors that one seeks to remove, and the gray subspace to be the space orthogonal to the frequency components one seeks to remove. The first projection moves the data into the space orthogonal to the motion components. Thus, the influence of motion has been effectively removed and the data, represented by the yellow point, is uncorrelated with motion. The second projection moves the data into the space orthogonal to the frequency components one seeks to remove, thereby removing the effects of these components. The data, represented by the green point, is in the frequency band of interest. If the spaces are orthogonal, as in the left plot, the data continue to be uncorrelated with motion and in the frequency band of interest, regardless of which order the projections are performed. However, if these two subspaces are not orthogonal to one another, the second projection actually moves the data back into the space spanned by the motion regressors. This has the unfortunate effect of reintroducing their effects into the data. Doing the projections in the reverse order would leave a point uncorrelated with the motion regressors, but not in the frequency band of interest.

### 2.2 Preprocessing steps as projections

While we have illustrated the problem in the context of motion regression and temporal filtering, we stress that it is not limited to these particular cases. The problem potentially arises in any setting when multiple preprocessing steps that act as linear projections are used sequentially. Preprocessing steps that fit this description include the removal of WM and CSF, CompCor, global signal regression, and spike regression.

In each of these steps, one can define a design matrix **X**_*i*_ consisting of the nuisance components one seeks to remove from the BOLD signal. These design matrices are then implicitly used to create a projection matrix

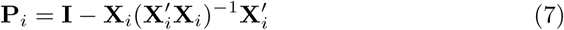

such as as the ones described above. The matrix is then used to project the data onto a subspace that is orthogonal to the space spanned by the nuisance components, thus removing their effect.

For example, filtering can be performed within a linear framework by including sine and cosine functions of the appropriate frequencies into the design matrix. Spike regression can be performed by including delta functions corresponding to each time point one seeks to remove into the design matrix. Nuisance regression and CompCor can be performed in a similar manner as outlined for motion regression.

### 2.3 Interaction among preprocessing steps

Each projection matrix **P**_*i*_, as defined in the previous section, projects the data onto an subspace that is orthogonal to the space spanned by the nuisance components included in the design matrix **X**_*i*_. As illustrated in Section 2.1, if a series of projections are applied sequentially there is a risk that the data is projected back into a subspace that one seeks to avoid.

Suppose we seek to perform *m* different modular preprocessing steps. The potential interaction between preprocessing steps can be explored by computing the product of the projection matrices **P**_1_**P**_2_ ⋯ **P**_*m*_ and evaluating whether or not it lies in the subspaces spanned by the columns of **X**_*i*_. Here, in order to not reintroduce nuisance signal contained in **X**_*i*_ we require

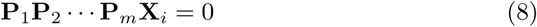

for all *i* = 1, *… m*.

If this condition does not hold, nuisance components will be reintroduced into the signal. Here the order in which the preprocessing steps are performed becomes important. Consider the following two preprocessing pipelines: (i) motion regression followed by temporal filtering; and (ii) temporal filtering followed by motion regression. In the first case motion is reintroduced by temporal filtering, while in the second case the filtered frequency components are reintroduced by motion regression. Which of these is more detrimental to the subsequent analysis can be debated, but regardless the reintroduction of unwanted nuisance components will ultimately change the interpretation of the findings.

As a general rule-of-thumb, the nuisance components related to the last preprocessing step performed should be adequately removed from the signal. This can be seen by noting that **P**_*m*_**X**_*m*_ = 0 is always true, and thus Eq. 8 holds. However, components corresponding to earlier steps are potentially reintroduced if not handled properly.

### 2.4 Recommendations

The problem of reintroducing nuisance regressors can be avoided in two ways. First one can define an omnibus projection matrix consisting of all nuisance variables one seeks to remove from the data. This entails performing motion regression, CompCor, temporal filtering, etc. in a common joint model. This is simple to do in a linear model framework by concatenating the *m* design matrices into a large omnibus design matrix:

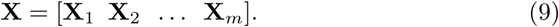

Using this design matrix ensures that the data is projected into a subspace that is orthogonal to all nuisance components contained in **X**.

Another approach is to create the design matrices in such a manner that they project the data onto a series of orthogonal subspaces. For example, spike regression projects the data onto a subspace orthogonal to the removed time points. If motion regression is subsequently performed then it is necessary to remove the same time points as in the spike regression from the motion design matrix. If not, the contribution of these time points are reintroduced into the signal. Another example is if temporal filtering is applied to the data before motion regression. Here both the data and the motion regressors must be filtered to avoid reintroducing the filtered bands back into the data after motion regression. This general point can be summarized as: one must orthogonalize a linear filter or set of nuisance covariates with respect to all previously removed sets of covariates to avoid re-introducing nuisance signals removed in previous steps.

## 3 Methods

While in Section 2 we focused on showing theoretically how the use of sequential modular preprocessing steps can reintroduce previously removed nuisance signal, in this section we focus on showing this empirically. We show this in the context of motion regression and high-pass filtering applied to test-retest rs-fMRI data. We explore how the use of modular preprocessing pipelines reintroduce artifacts that have been previously removed. We also seek to show that using a joint (non-modular) pipeline circumvents these issues.

### 3.1 Data collection

We used the Multi-Modal MRI Reproducibility Resource (Kirby) from the F.M. Kirby Research Center^2^; see Landman et al.(2011) for a detailed explanation of the acquisition protocol. Briefly, it consists of data from 21 healthy adults scanned on a 3T Philips Achieva scanner. Each participant completed two scanning sessions on the same day. Between sessions the participants briefly exited the scan room and a full repositioning of the participant, coils, blankets, and pads occurred prior to the second session. A T1-weighted MPRAGE structural run was acquired during both sessions (acquisition time = 6 *min*, TR/TE/TI = 6.7*/*3.1*/*842 *ms*, resolution = 1×1×1.2 *mm*^3^, SENSE factor = 2, flip angle = 8°). A multi-slice SENSE-EPI pulse sequence (Stehling et al., 1991;Pruessmann et al.,1999) was used to acquire one rs-fMRI run during each session, where each run consisted of 210 volumes sampled every 2 *s* at 3 *mm* isotropic spatial resolution (acquisition time: 7 *min*, TE = 30 *ms*, SENSE acceleration factor = 2, flip angle = 75°, 37 axial slices collected sequentially with a 1 *mm* gap). Participants were instructed to rest comfortably while remaining as still as possible, and no other instruction was provided. We will refer to the first rs-fMRI run collected as Session 1 and the second as Session 2. One participant was excluded from data analyses due to excessive motion.

### 3.2 Initial preprocessing

SPM 12 (Wellcome Trust Centre for Neuroimaging, London, United Kingdom) and custom MATLAB (The Mathworks, Inc., Natick, MA) scripts were used to preprocess the Kirby data. To allow for the stabilization of magnetization, four volumes were discarded at acquisition, and an additional volume was discarded prior to preprocessing. Slice timing correction was performed using the slice acquired at the middle of the TR as reference, and rigid body realignment parameters were estimated to adjust for head motion. Structural runs were registered to the first functional frame and spatially normalized to Montreal Neurological Institute (MNI) space using SPM’s unified segmentation-normalization algorithm (Ashburner and Friston,2005). The estimated rigid body and nonlinear spatial transformations were applied to the rs-fMRI data.

### 3.3 Creation of motion images

Before proceeding with further preprocessing steps, we estimated the motion at each voxel of the brain. To do so we used a design matrix **X** consisting of 24 regressors. These included the six motion regressors obtained after rigid-body transformation, the regressors squared, their first order difference, and the first order difference squared. After estimating the parameters *β*_*v*_ corresponding to these regressors at each voxel *v*, we proceeded to create ‘motion images’. This was done by computing the fitted values **X***β*_*v*_ at each voxel. Thus, we were able to create 4D images of the estimated contribution of motion to the signal at each voxel in the brain. Ultimately, the goal of preprocessing is to remove the effects of motion. However, at this stage we do not remove this component, but simply compute it to use as a baseline to evaluate the motion related components left in the signal after performing specific preprocessing steps.

### 3.4 Secondary preprocessing pipelines

In the next stage of preprocessing we used three different preprocessing pipelines. In the first pipeline, we first perform motion regression (as described above) and remove the estimated motion from the data. Thereafter we apply a high pass filtered using a cutoff frequency of 0.01 *Hz*. We refer to this pipeline as ‘MCHP’. In the second pipeline, we begin by applying the high pass filter, and thereafter perform motion regression to remove the estimated motion from the data. We refer to this pipeline as ‘HPMC’. Finally, in the third pipeline we perform motion regression and high-pass filtering jointly using a combined design matrix. We refer to this pipeline as ‘Joint’.

### 3.5 Evaluation of pipelines

Next we sought to evaluate the interaction between motion regression and highpass filtering in each pipeline. For each pipeline and after each modular step, we computed the correlation between the preprocessed data and the motion images at each voxel. This was done to evaluate the residual contribution of motion after performing each preprocessing step.

We also parcellated the data at each stage into 268 regions using the Shen atlas (Shen et al.,2013), and computed the correlation matrices across regions as commonly done in rs-fMRI studies on whole brain functional connectivity. This was repeated for both sessions for each of the 20 subjects. For each session and pipeline we computed the average correlation matrices at the group-level. We also performed t-tests to determine whether the Fisher-transformed correlation between the motion time courses and the data after motion regression followed by high-pass filtering was significantly different from 0.

In addition, for each pipeline, after each step we estimated the spectral density for every voxel time course in order to evaluate the relative contribution of different frequencies components. In particular, we focused on the contribution of frequencies lower than 0.01 *Hz*, as these are the ones we sought to remove from the signal. We will quantify this contribution by computing the proportion of the power that lies in the frequency band below 0.01 *Hz* in each voxel. This is a measure that is similar in spirit to fractional Amplitude of Low Frequency Fluctuations (fALFF; Zou et al.(2008))

## 4 Results

To properly understand the effects of modular preprocessing pipelines we illustrate its effects on a single voxel, a single subject, and group analysis.

### 4.1 Illustration of voxel-level effects

Figure 4 illustrates the problem at a single voxel of the brain. It shows an example fMRI time course and the estimated motion time course, which are highly correlated with one another (*r* = 0.7622); see Panels A and B. Here A shows plots of the two time courses, while B shows the relationship using a scatter plot. After motion regression the resulting times series (Panel C) is uncorrelated with the motion (Panel D), which is to be expected. However, high-pass filtering has the effect of reintroducing the correlation between the data and the motion, which is now *r* =*-*0.3458 (Panels E-F). This illustrates how the two preprocessing steps interact with one another. The high-pass filter has the effect of projecting the data back into the space spanned by the motion regressors, thus reintroducing a correlation between the signal and the motion. Note that if high-pass filtering is performed before motion correction, the resulting time course is uncorrelated with the motion (Panels G-H). However, this leads to the reintroduction of frequency components that had previously been removed. This can clearly be seen studying the periodograms shown in Figure 5. Prior to the secondary preprocessing steps there is clear signal in the range below 0.01 *Hz*; see Panel A. If motion correction is performed followed by high-pass filtering than signal in this range disappears as seen in Panel B. However, if the order of these preprocessing steps are reversed it is clear from Panel C that signal is reintroduced into these frequencies.

**Figure 4:**
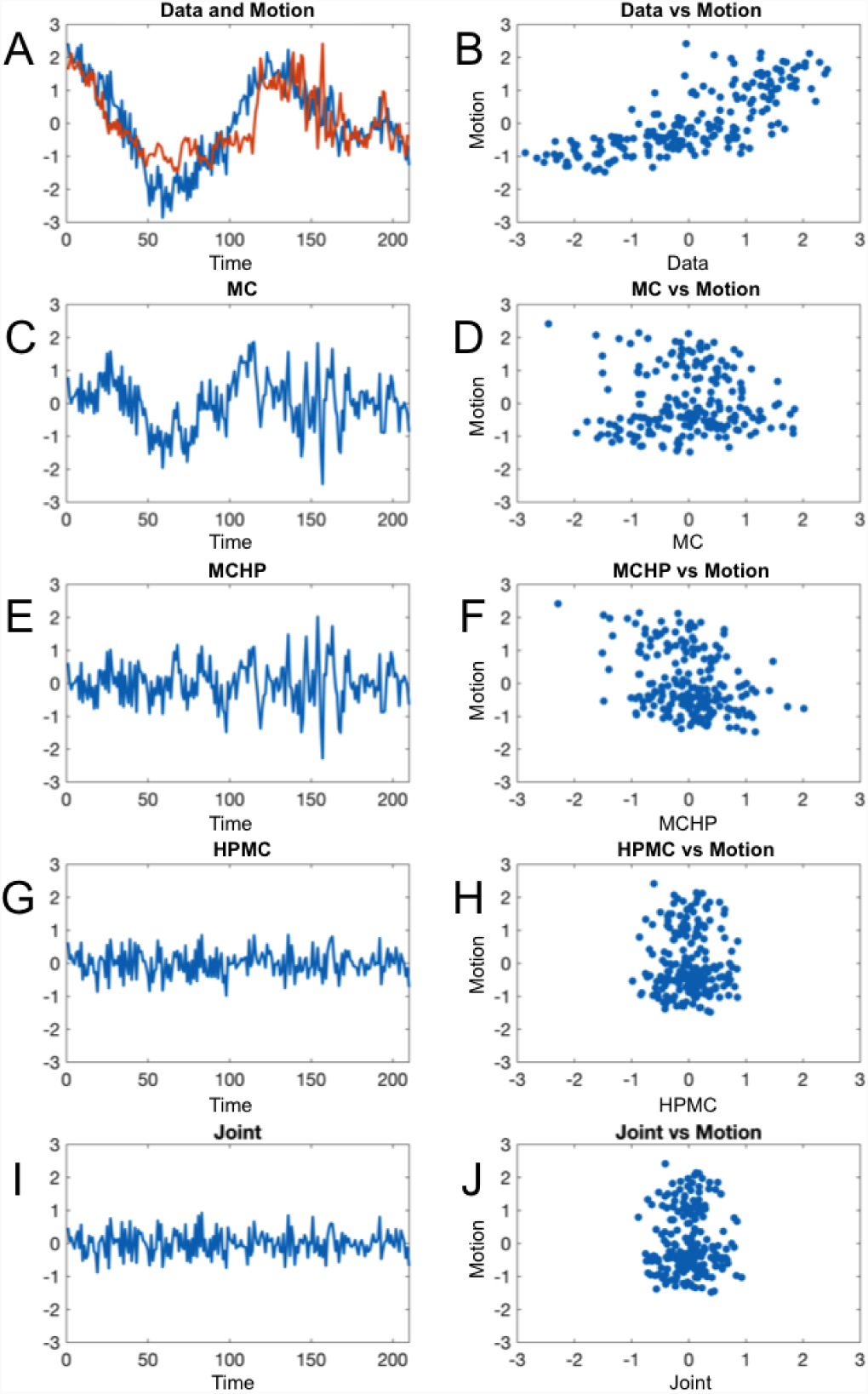
Results from an example voxel. (A) The rs-fMRI time course before secondary preprocessing (blue) and the estimated motion (red) from the voxel of interest. (B) A scatter plot of these two time courses show a positive correlation (*r* = 0.7622). (C) The data after motion correction. (D) A scatter plot of this time course and the estimated motion show no correlation (*r* = 0). (E) The data after motion correction fooled by high-pass filtering. (F) A scatter plot of this time course and the estimated motion show a negative correlation (*r* = *-*0.3458). (G) The data after high-pass filtering followed by motion correction. (H) A scatter plot of the time course and the estimated motion show no correlation (*r* = 0). (I) The data after joint high-pass filtering and motion correction. (J) A scatter plot of the the time course and the estimated motion show no correlation (*r* = 0)

**Figure 5:**
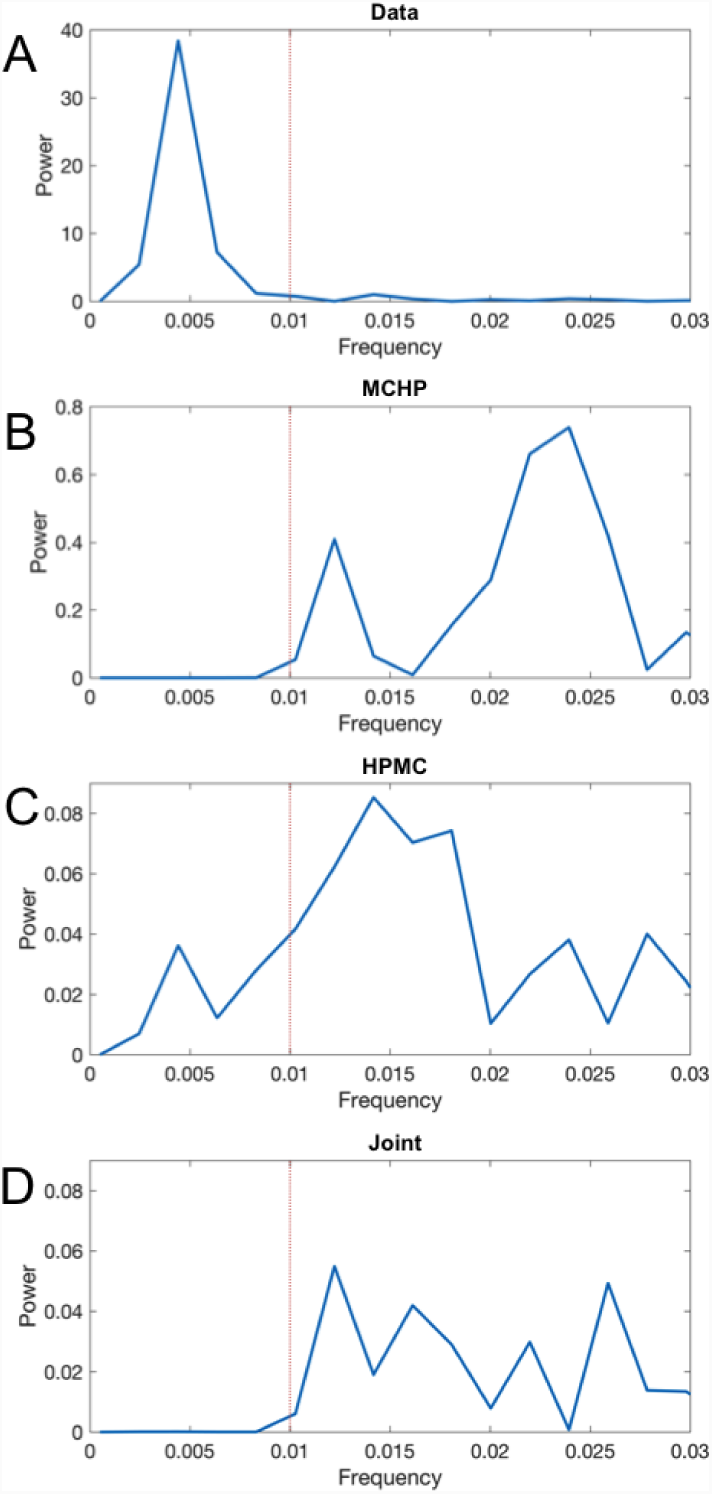
Results from an example voxel. (A) The periodogram in the range [0 0.03] *Hz* of the rs-fMRI time course before secondary preprocessing. (B) The periodogram of the data after motion correction and high-pass filtering. (C) The periodogram after high-pass filtering and motion correction. (D) The periodogram after joint high-pass filtering and motion correction. In each panel the red dotted line reflects the frequency cut-off of the high-pass filter.

One approach towards handling this problem is to perform both operations within a joint model. As can be seen from Figures 4I-J and 5D, performing joint motion correction/temporal filtering provides a time course that is uncorrelated with motion, while only containing information in the frequencies of interest. Thus, in this approach the effects of both motion regression and temporal filtering are retained in the processed signal as intended.

### 4.2 Illustration of whole-brain effects

The point illustrated above is reinforced by looking at each voxel from a randomly chosen subject; see Figure 6. First we focus on studying the reintroduction of motion. Panel A shows the correlation between the motion regressed data and the motion at each voxel of the brain. Clearly the correlation is negligible (< 2 × 10^−6^) across all voxels. However, after high-pass filtering the correlation is reintroduced; see Panel B. Interestingly, the correlation between the data and motion is now mostly negative. Panel C shows these correlations superimposed onto the brain. Clearly, there are patterns consistent with motion artifacts, including a ring-like shape at the edge of the brain. There are also apparent patterns in the frontal cortex.

**Figure 6:**
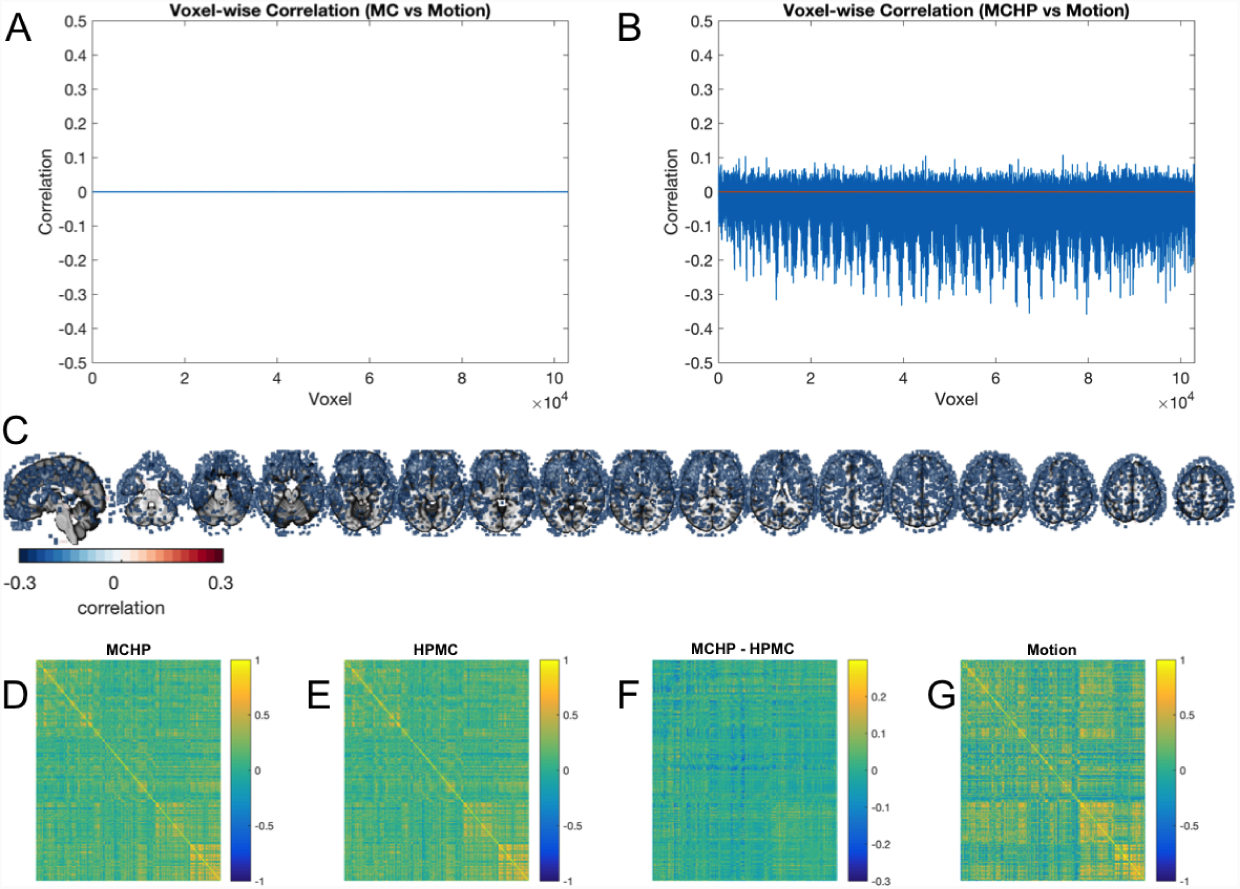
(A) The correlation between the motion regressed data and the motion at each voxel of the brain is negligible (< 2 ×10^−6^). (B) The correlation is reintroduced after high-pass filtering. (C) Correlations from (B) superimposed onto their brain locations. They are threholded at an arbitrary value OF of ± 0.05. (D) The correlation matrix between the 268 regions of the Shen atlas computed on data where motion regression is performed followed by high-pass filtering. (E) The same correlation matrix on data where high-pass filtering is performed followed by motion regression. (F) The difference between the correlation matrices in E and F. (G) The correlation matrix between the motion time courses over the same regions.

To further illustrate the impact of the reintroduction of motion we study the correlation matrices across the 268 regions of the Shen atlas (Shen et al., 2013) both when (i) performing motion regression followed by high-pass filtering; and (ii) performing high-pass filtering followed by motion regression. In the first case the motion is reintroduced by temporal filtering, while in the second case it should be removed. Note that in the second case, filtered frequency components are instead reintroduced (see Section 2.3). Panels D and E show the results, and F shows the difference between the two correlation matrices. Clearly the differences are substantial as it ranges between values of ±0.3. Interestingly, the patterns seen in the difference (Panel F) shows similarities to the correlation between the motion time courses over the same regions (Panel G).

Note that if high-pass filtering is performed before motion correction, we deal with a different problem as illustrated in Figure 7. Panel A shows the proportion of the power that lies in the frequency band below 0.1 *Hz* in each voxel after motion correction followed by high-pass filtering. Here we can note that as expected there is a negligible contribution as all values are < 3 × 10^−3^. However, Panel B shows the same plot for data where motion correction is performed after high-pass filtering. The plot shows there has been a significant reintroduction of information from frequencies that had previously been removed. In certain voxels up to 12% of the power lies in frequencies that should ideally be 0. The spatial position of these voxels are shown in Panel D, where the results are superimposed onto the brain. The results are presented using an arbitrary threshold of 2% to better be able to identify voxels with a high proportion of power in the low frequency band. Finally, Panel C shows the same plot for data preprocessed jointly. Much like in Panel A there a negligible contribution of signal in frequencies below 0.01 *Hz*.

**Figure 7:**
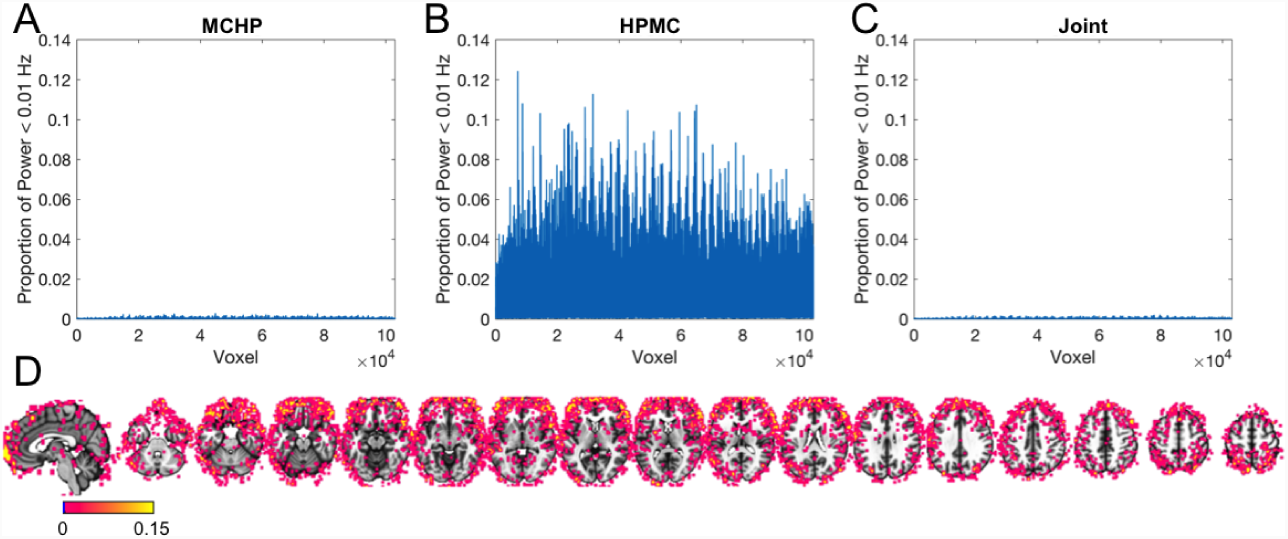
(A) The proportion of the power that lies in the frequency band below 0.01 *Hz* in each voxel after motion correction followed by high-pass filtering. Note there is a negligible contribution. (B) Same plot for data where motion correction is performed after high-pass filtering. In certain voxels up to 12% of the power lies in frequencies that should ideally be 0. (C) Same plot for data processed jointly. Much like in (A) there a negligible contribution. (D) The results shown in (B) superimposed onto the brain. Note the results are presented using an arbitrary threshold of 2% for visualization purposes.

### 4.3 Illustration of group-level effects

Finally, we turn our attention to group level analysis. Here we analyze the two sessions of the Kirby data set separately. The results related to motion are shown in Figure 8. Panels A-C show the results from the first session. Panels A and B show the average correlation matrices when performing motion regression followed by high-pass filtering and performing high-pass filtering followed by motion regression, respectively. Panel C shows the difference between these two correlation matrices. Panels D-F show equivalent results for Session 2. In both cases the differences range between values of ±0.06. This illustrates that while averaging across subjects has removed some of the differences between preprocessing streams there are still substantial differences that remain, and are consistent in sign across participants, introducing systematic bias in group-level results.

**Figure 8:**
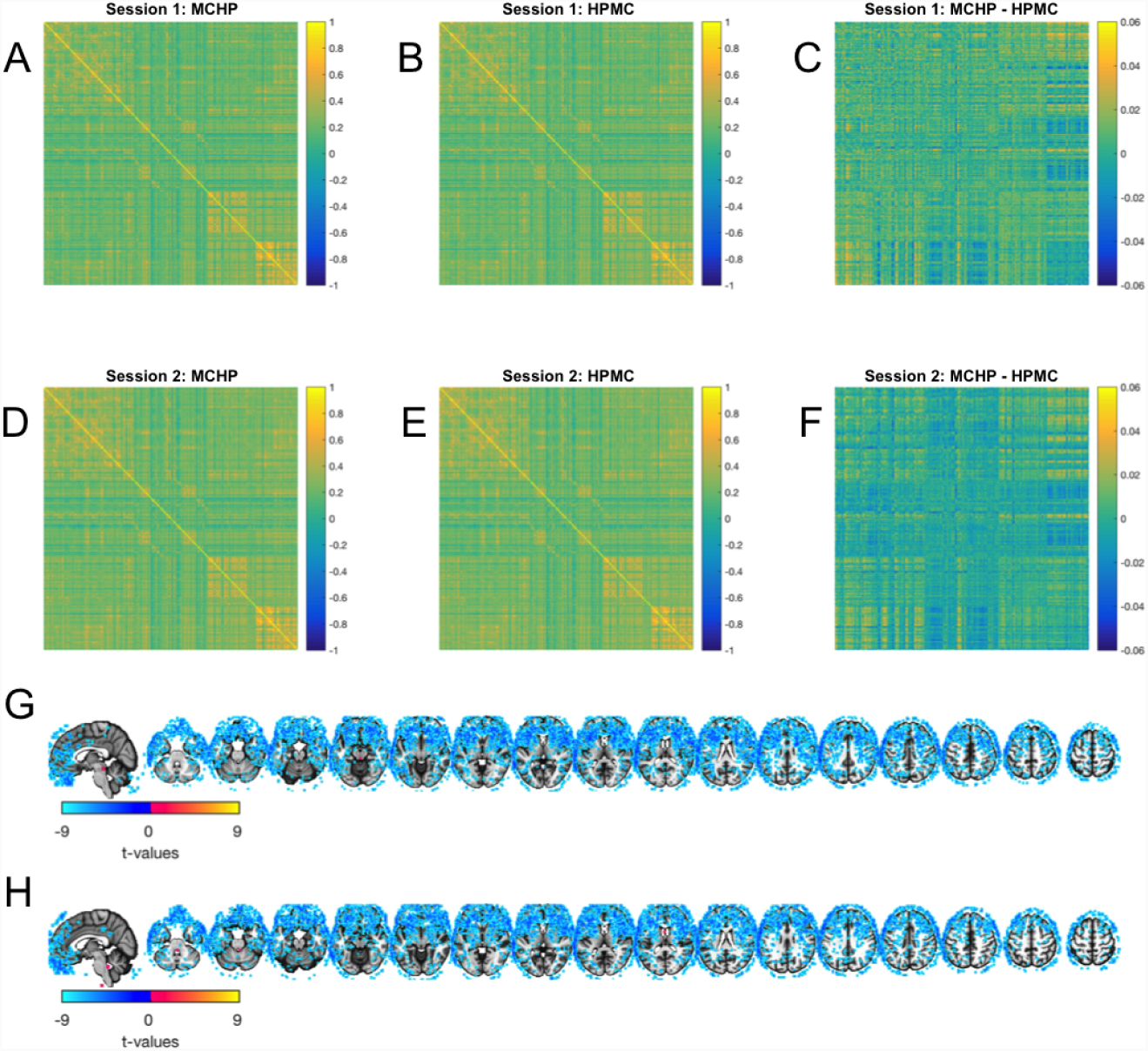
(A) The average correlation matrix when performing motion regression followed by high-pass filtering for the 20 subjects in Session 1. (B) The average correlation matrix when performing high-pass filtering followed by motion regression for the 20 subjects in Session 1. (C) The difference between the two correlation matrices shown in (A) and (B). (D)-(F) Equivalent results for Session 2. (G) Group-level t-maps for Session 1 for determining whether the Fisher-transformed correlation correlation between the motion time courses and the data after motion regression followed by high-pass filtering is significantly different from 0. Results thresholded at *p* < 0.001. (H) Same results for Session 2.

Panels G and H show the results from each session of t-tests to determine whether the Fisher-transformed correlation correlation between the motion time courses and the data after motion regression followed by high-pass filtering is significantly different from 0. Clearly, the results are very similar across sessions and there is clear correlation in frontal areas of the brain.

Figure 9 shows the group average proportion of the power that lies in the frequency band below 0.01 *Hz* at each voxel for both sessions using the high-pass filtering followed by motion regression pipeline. There are clear similarities between the results between the two sessions. As seen in previous figures there are patterns consistent with motion artifacts, including a ring-like shape at the edge of the brain. There are also apparent patterns in the frontal cortex.

**Figure 9:**
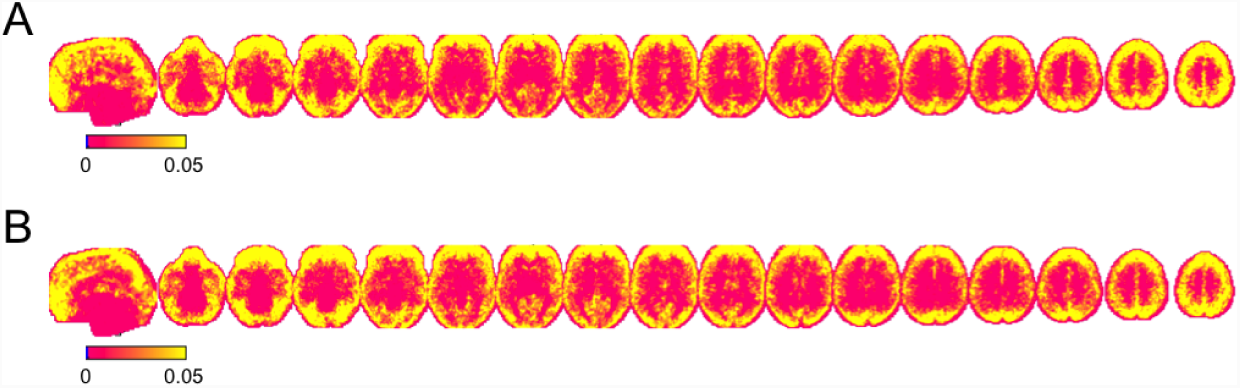
(A) Group-level averages of the proportion of the power that lies in the frequency band below 0.1 *Hz* at each voxel for Session 1. (B) Same results for Session 2.

## 5 Discussion

It is common practice in the field of rs-fMRI for researchers to piece together modular preprocessing pipelines consisting of a number of separately developed algorithms; each designed to remove a specific type or class of artifacts. These artifacts can be related to scanner drift, signal spikes, motion, and signal fluctuations due to heart rate and respiration. In this paper we offer a critic of this modular preprocessing approach, and argue that using such a pipeline can potentially have adverse effects on subsequent statistical analysis (e.g., restingstate functional connectivity).

While all preprocessing steps performed on fMRI data are important, we argue there needs to be a clear understanding about the effects they have on both the spatial and temporal correlation structure. More generally, it is critical to study the interactions among the individual preprocessing steps. In this work we have proposed a geometrical framework that provides a way to better understand these interactions. The framework can be used to evaluate any preprocessing step that can be expressed within a linear model framework; examples of such include motion regression, scrubbing, removal of WM and CSF signal, global signal removal, and temporal filtering. Together these account for a large segment of the set of possible preprocessing steps applied to rs-fMRI data.

Using our approach we were able to illustrate how preprocessing steps performed at a later stage of the pipeline can potentially reintroduce artifacts that had previously been removed from the data in an earlier step. Potential interactions between preprocessing steps are explored by computing the product of their respective projection matrices and evaluating whether or not when applied together they project the data into a subspace spanned by the various nuisance components. If this is the case, these nuisance components will be effectively reintroduced into the signal in a manner consistent with the order in which they were performed. Hence, the order in which the steps are organized in the preprocessing pipeline is critical. As a rule-of-thumb, the nuisance components corresponding to the last preprocessing step performed should be adequately removed from the signal, with components corresponding to earlier steps potentially reintroduced if they are not orthogonal to components removed in subsequent steps. We illustrate these issues both theoretically and using test-retest fMRI data. Empirically we find that the reintroduced artifacts are consistent across sessions, and can potentially influence findings at both the single subject and group level.

We note that the issues discussed in this paper can be circumvented in two ways. The first approach would be to abandon the modular approach to preprocessing the data, and instead use a joint approach that simultaneously performs the different preprocessing steps within an omnibus framework. For example, it would be relatively straightforward to formulate a single linear model that simultaneously performs motion regression, nuisance regression, and temporal filtering. This is an approach advocated by Caballero-Gaudes and Reynolds (2017). We agree that the development of models that incorporate multiple preprocessing steps promises to play an important role in the future (Lindquist et al., 2008). The second approach is to formulate the design matrices used in each preprocessing step in such a manner that when applied sequentially they are constrained to project onto orthogonal subspaces. For example, if motion correction is performed after temporal filtering, the columns of the design matrix used in the motion regression should also be temporally filtered in a similar manner to ensure the data is projected onto an orthogonal subspace. The general rule-of-thumb is that one must orthogonalize both the data, and all subsequent projections to maintain data orthogonality with the current projection.

Interestingly, these approaches are often utilized when analyzing task fMRI using the general linear model (GLM). Here it is common to perform motion regression and temporal filtering simultaneously by including both terms in the design matrix. If these steps are instead performed modularly (i.e., with low-pass filtering performed prior to fitting the GLM) the same issues described in this paper will impact the results unless the design matrix used in the GLM has also been temporally filtered. Taking this step ensures that the frequencies removed in the low-pass filtering are not reintroduced when fitting the GLM. This is actually the default in most common fMRI software packages (e.g., SPM and FSL). Hence, it is not clear if the issues raised in this paper are problematic for most task fMRI studies, and therefore we focus on rs-fMRI, though we urge researchers to be cautious when performing modular analysis to ensure that artifacts are not reintroduced.

The complete preprocessing of rs-fMRI data can conveniently be separated into spatial and temporal pre-processing pipelines (Smith et al., 2013). The goal of spatial pre-processing is to remove spatial artifacts from the data. Here the relevant steps include correction for spatial distortions caused by gradient nonlinearity, rigid-body correction for motion, correction for *B*_0_ distortion, co-registration of structural and functional data, and normalization to a standard template (i.e., MNI-space). These steps are performed prior to temporal pre-processing, which includes motion regression, scrubbing, removal of WM and CSF signal, global signal removal, and temporal filtering. In this work we have focused entirely on potential problems with temporal preprocessing, and our geometrical framework is not explicitly designed for evaluating spatial preprocessing pipelines. Future work involves extending the framework, or alternatively developing a companion framework, for working in this setting.

It appears to be a trend for large data consortium (e.g., the Human Connectome Project) to make minimally preprocessed data available (Glasser et al., 2013). These data have only undergone spatial preprocessing, and should thus not be effected by the problems outlined in this paper. However, we note that once researchers take these data sets and perform temporal preprocessing, they are susceptible to the problems outlined herein.

It is difficult to provide an exact estimate of the number of rs-fMRI papers that suffer from the issues raised in this work. This is in large part due to the wide variability in the reporting of preprocessing methods used in rs-fMRI studies (Waheed et al., 2016). In general, methods sections are often extremely terse when discussing preprocessing, and neither specify the order or the manner in which the different steps are performed. In addition, even if the order is specified it is not always clear whether proper orthogonalization has been performed. That said we suspect that the problems outlined in this paper negatively impact a large number of papers published every year.

## 6 Acknowledgements

The work presented in this paper was supported in part by NIH grants R01 EB016061 and R01 EB026549 from the National Institute of Biomedical Imaging and Bioengineering.

## Footnotes

1 Note this survey also included task-based fMRI studies.

2 Publicly available at http://www.nitrc.org/projects/multimodal

